# Identification of a small chemical as a lysosomal calcium mobilizer and characterization of its ability to inhibit autophagy and viral infection

**DOI:** 10.1101/2022.06.09.495506

**Authors:** Kehui Zhang, Lihong Huang, Nanjun Chen, Jianbo Yue

## Abstract

We previously identified GADPH as one of the cyclic adenosine diphosphoribose (cADPR)’s binding proteins and found that GADPH participates in cADPR-mediated Ca^2+^ release from ER via RyRs. Based on the simulated cADPR-GAPDH complex structure, we performed the structure-based drug screening, identified several small chemicals with high docking scores to cADPR’s binding pocket in GAPDH, and showed that two of these compounds, C244 and C346, are potential cADPR antagonists. We further synthesized several analogs of C346, and found that its analog, G42, also mobilized Ca^2+^ release from lysosomes. G42 alkalized lysosomal pH, and inhibited autophagosome-lysosome fusion. Moreover, G42 markedly inhibited Zika virus (ZIKV, a flavivirus) or murine hepatitis virus (MHV, a β-coronavirus) infections of host cells. These results suggest that G42 inhibits virus infection, likely by triggering lysosomal Ca^2+^ mobilization and inhibiting autophagy.

## INTRODUCTION

Cyclic adenosine diphosphoribose (cADPR), an endogenous Ca^2+^ mobilizing nucleotide, elicits Ca^2+^ release from endoplasmic reticulum (ER). Its mediated Ca^2+^ signaling regulates diverse cellular processes, e.g., the circadian clock in plants and long-term synaptic depression in the hippocampus (Deterre et al., 2000; Galione and Churchill, 2000; Guse, 2004; Malavasi et al., 2008; Wei et al., 2014). cADPR targets ryanodine receptors (RyRs) on the ER in many cell types, yet cADPR does not act directly on RyRs. We thus synthesized a photoaffinity labeling cADPR analog (PAL-cIDPRE), applied it to the mammalian cell extracts, and identified GAPDH as a cADPR binding protein. We characterized cADPR’s binding affinity with GAPDH and mapped its binding sites in GAPDH. We further demonstrated that cADPR induces the interaction between GAPDH and RyRs, and GAPDH knockdown inhibited cADPR-mediated Ca^2+^ mobilization. Our results suggest that GAPDH is a cADPR binding protein and participates in cADPR-induced Ca^2+^ release from ER (Zhang et al., 2017).

In glycolysis, GAPDH catalyzes D-glyceraldehyde 3-phosphate to 1,3-bisphospho-D-glycerate with NAD^+^ as co-enzyme, in which NAD^+^ is reduced to NADH (Seidler, 2013). Although traditionally viewed as a “housekeeping gene”, GAPDH is a multifunctional protein, playing key roles ranging from metabolism, autophagy, apoptosis, mRNA stability, intracellular membrane trafficking, heme metabolism, genomic integrity, and nuclear tRNA export (Chang et al., 2015; Chang et al., 2013; Yun et al., 2015). Extra- or intracellular stress could modulate distinct pools of GAPDH via the dynamic changes of GAPDH’s subcellular localization, or/and posttranslational modifications, or/and protein-protein interactions for its multifunctional roles, thereby enabling cells to adapt to stresses (Hara et al., 2005; Sawa et al., 1997; Tristan et al., 2011). Dysregulated GAPDH might lead to numerous human diseases, e.g., neurodegenerative disease and heart failure (You et al., 2013).

The Ca^2+^ signaling participates in many steps of the virus infection cycle, e.g., entry, replication, packing, and egress. Manipulation of Ca^2+^ signaling by small chemicals provides a therapeutic option to control virus infection (Alam et al., 2016; Bissig et al., 2013; Fujioka et al., 2018; Fujioka et al., 2013; Haughey and Mattson, 2002; Sakurai et al., 2015; Scherbik and Brinton, 2010; Wang et al., 2017; Zhou et al., 2009). Here, we performed structure-based drug screening and identified several cADPR analogs, such as C346, based on the simulated cADPR-GAPDH complex structure. We then synthesized several analogs of C346, and further characterized the Ca^2+^ mobilization, autophagy modulation, and antiviral activity of one of its analogs, G42. We demonstrate that G42 inhibits virus infection, likely by mobilizing lysosomal Ca^2+^ release and inhibiting autophagy.

## RESULTS and DISCUSSION

### Virtual screening and pharmacological characterization of novel cADPR agonist(s) or antagonist(s)

Based on the simulated GAPDH-cADPR complex structure (Zhang et al., 2017), we performed the virtual drug screening to identify potential cADPR agonist(s) or antagonist(s). Briefly, compounds in the ZINC library were pretreated and minimized in Discovery Studio 2016 (DS 2016), and subsequently docked to cADPR’s binding pocket in GAPDH by the GOLD docking protocol. Afterward, the list of docking scores showing the binding affinity of each compound to GAPDH was generated. The top one thousand molecules with potential bioactivity were further screened and filtered manually (**Fig. 1A**). Finally, ten potential small chemicals with high docking scores to cADPR’s binding pocket in GAPDH were identified (**Fig. 1B**). One of them, compound 244 (C244), is commercially available, and we found that C244 itself did not affect cytoplasmic Ca^2+^ levels (**Fig. 1C**), but pretreatment of Jurkat cells (**Fig. 1D**) or RyR3-expressing HEK293 cells (**Fig. 1E**) with C244 abrogated NPE-cADPR-mediated Ca^2+^ release, suggesting that C244 is a potential cADPR antagonist. Similarly, we found that another compound, 346 (C346), by itself did not affect cytoplasmic Ca^2+^ levels (**Fig. 1F**), but pretreatment of Jurkat cells (**Fig. 1H**) or RyR3-expressing HEK293 cells (**Fig. 1G**) with C346 inhibited NPE-cADPR or cIDPRE-mediated Ca^2+^ release, suggesting that C346 is also a potential cADPR antagonist.

**Figure 1.**
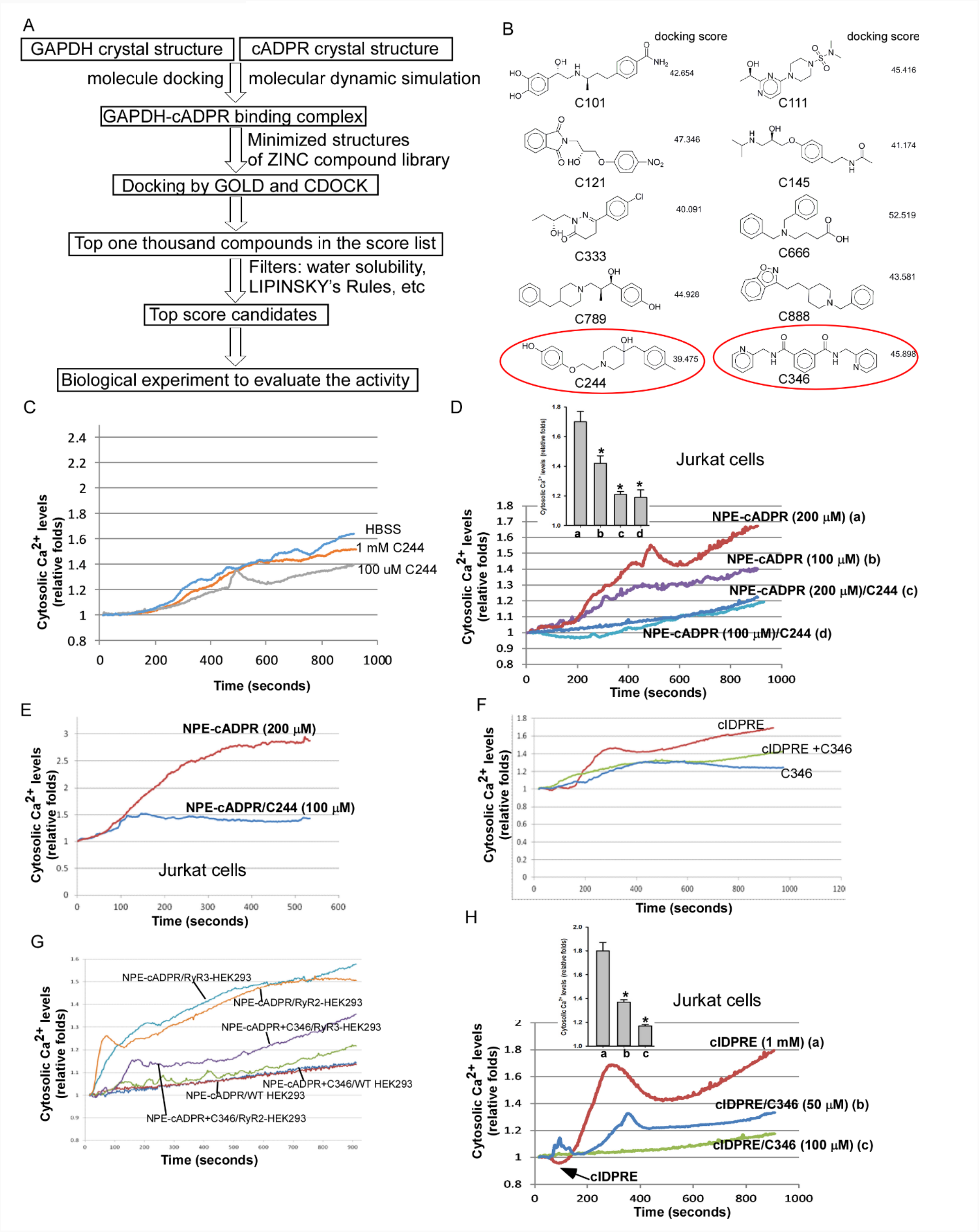
Identification of cADPR antagonists by structure-based drug screening. (**A**) Schematic diagram of drug screening. (**B**) 10 compounds with high docking scores were identified. (**C-E**) C244 failed to induce Ca^2+^ release (**C**), but pretreatment of Jurkat cells (**D**) or RyR3-expressing HEK293 cells (**E**) with 100 μM or 50 μM of C244 significantly inhibited NPE-cADPR (100 μM)-induced cytosolic Ca^2+^ increase. (**F-H**) C346 failed to induce Ca^2+^ release (**F**), but pretreatment of Jurkat cells (**G**) or RyR3-expressing HEK293 cells (**H**) with 100 μM or 50 μM of C346 significantly inhibited NPE-cADPR (100 μM) or cIDPRE (1 mM))-induced increase in cytosolic Ca^2+^.

Next, we performed surface plasmon resonance (SPR) experiments and found that C244 or C346 bound to the recombinant GAPDH protein immobilized on the CM5 chip (**Figs 2A** and **2B**). We then analyzed the binding mode of cADPR, C244, or C346 to GAPDH using the Schrödinger software (**Fig. 2C**). We previously showed that the His179 and Arg234 residues of GAPDH are two key amino acid residues for binding to cADPR, which binds to GAPDH by forming a hydrogen bond with the pyrophosphate moiety (left panel of **Fig. 2C**) (Zhang et al., 2017). Compared to cADPR binding with GAPDH, C346 exhibited a distorted conformation due to the presence of two substituents at the meta position of the benzene ring, and its pyridine moiety can form a π–π stack with His179 of GAPDH (middle panel of **Fig. 2C**). Interestingly, C244 exhibited a stretched conformation and formed hydrogen bonds with amino acids on the other side of the pocket, thus occupying part of the cADPR binding site. In this way, it might hinder the entry of cADPR, accounting for its antagonistic activity (right panel of **Fig. 2C**).

**Figure 2.**
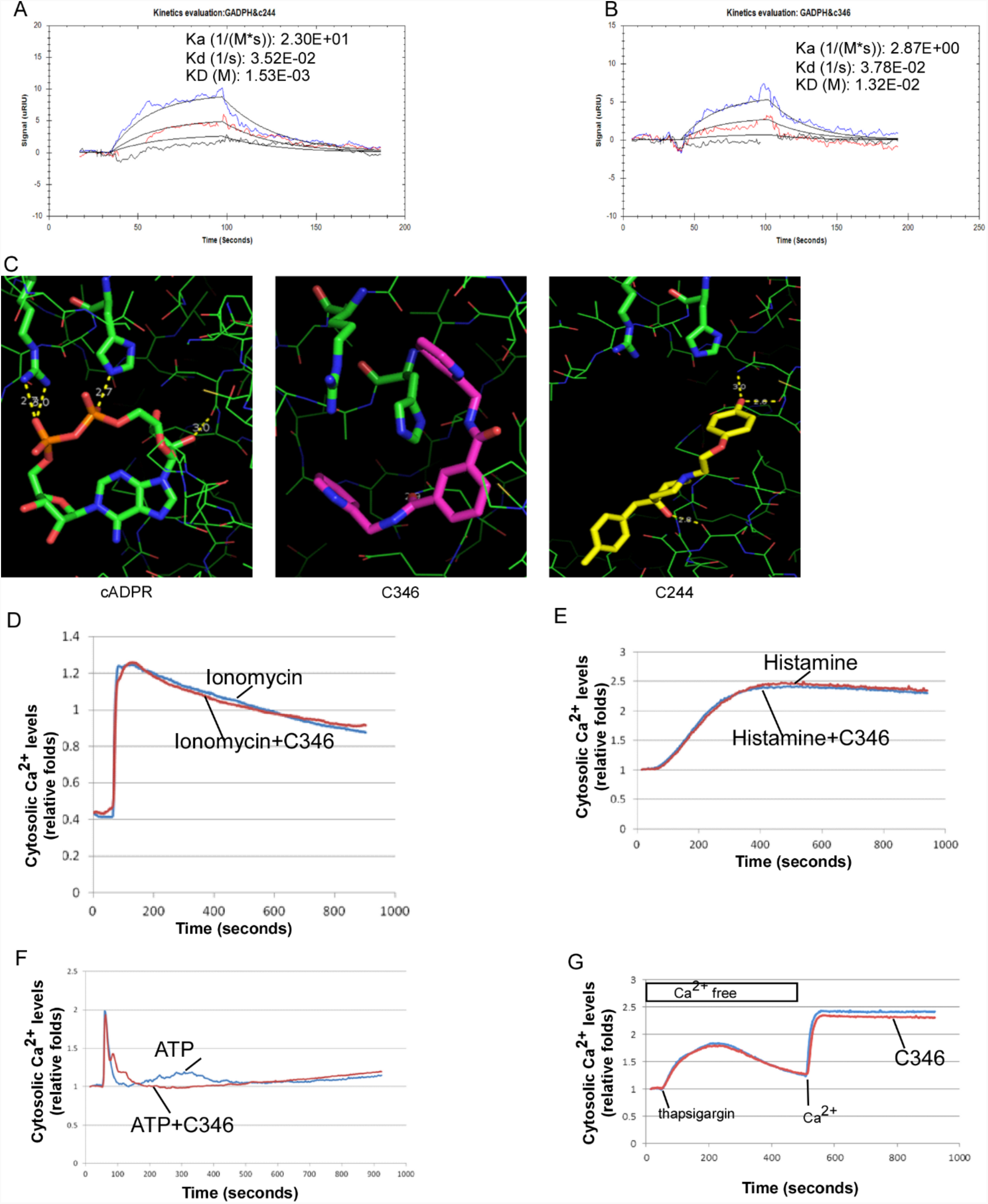
C346 is a cADPR-specific antagonist. (**A, B**) The affinity of C244 (**A**) or C346 (**B**) for GAPDH was determined by surface plasmon resonance (SPR). (**C**) The binding mode of GAPDH to cADPR, C244, or C346 was analyzed by Schrödinger software. (**D-G**) C346 (100 μM) pretreatment of HeLa cells did not significantly inhibit ionomycin (**D**), histamine (**E**), and ATP (**F**)-mediated Ca^2+^ release, nor did it affect store-operated Ca^2+^ influx (**G**).

C346 is a weak cADPR antagonist (**Figs 1G, 1H**), and we speculate that this might be due to the small molecular weight of C346, which cannot fully occupy the binding pocket of GAPDH (middle panel of **Fig. 2C**). Meanwhile, we found that C346 did not affect Ca^2+^ release mediated by ionomycin (**Fig. 2D**), histamine (**Fig. 2E**), and ATP (**Fig. 2F**), nor did it affect store-operated Ca^2+^ influx (**Fig. 2G**). These data suggest that C346 is a cADPR-specific antagonist.

### Chemical synthesis of C346 analogs and characterization of their Ca^2+^ mobilization activity

Subsequently, we set out to structurally modify C346, such as adding new pharmacophores to obtain larger molecules in order to fully occupy the cADPR binding pocket in GAPDH, or replacing functional groups of C346 to form new interactions with the amino acid residues in order to enhance the binding affinity (**Fig. 3A**). We synthesized more than 20 analogs of C346, but most of them were poorly soluble and were only soluble in DMSO. When they were added to the 24-well plate for testing, they precipitated almost immediately; thus, their activity could not be tested. We, therefore, only assessed six soluble compounds (G22, G31, G32, G41, G42, and G43) for their ability to modulate cytosolic Ca^2+^ concentration. To our surprise, in RyR3-expressing HEK293 cells, several of these compounds (50 μM) significantly triggered intracellular Ca^2+^ mobilization (**Fig. 3B**). However, compounds G22, G31, G41, G43, and G32 also significantly triggered intracellular Ca^2+^ mobilization in control HEK293 cells (lacking RyR expression) (**Figs 3C, 3D, 3F, 3H**, and **3I**). Only compound G32 or G42 triggered stronger intracellular Ca^2+^ mobilization in RyR3-expressing HEK293 cells than in wild-type HEK293 cells (**Figs 3E** and **3G**). Notably, G32 or G42 elicited a similar first peak of intracellular Ca^2+^ mobilization in RyR3-HEK293 cells when compared to control HEK293 cells (**Figs 3E** and **3G**). These results suggest that the second peak of intracellular Ca^2+^ mobilization triggered by G32 or G42 is RyRs-dependent.

**Figure 3.**
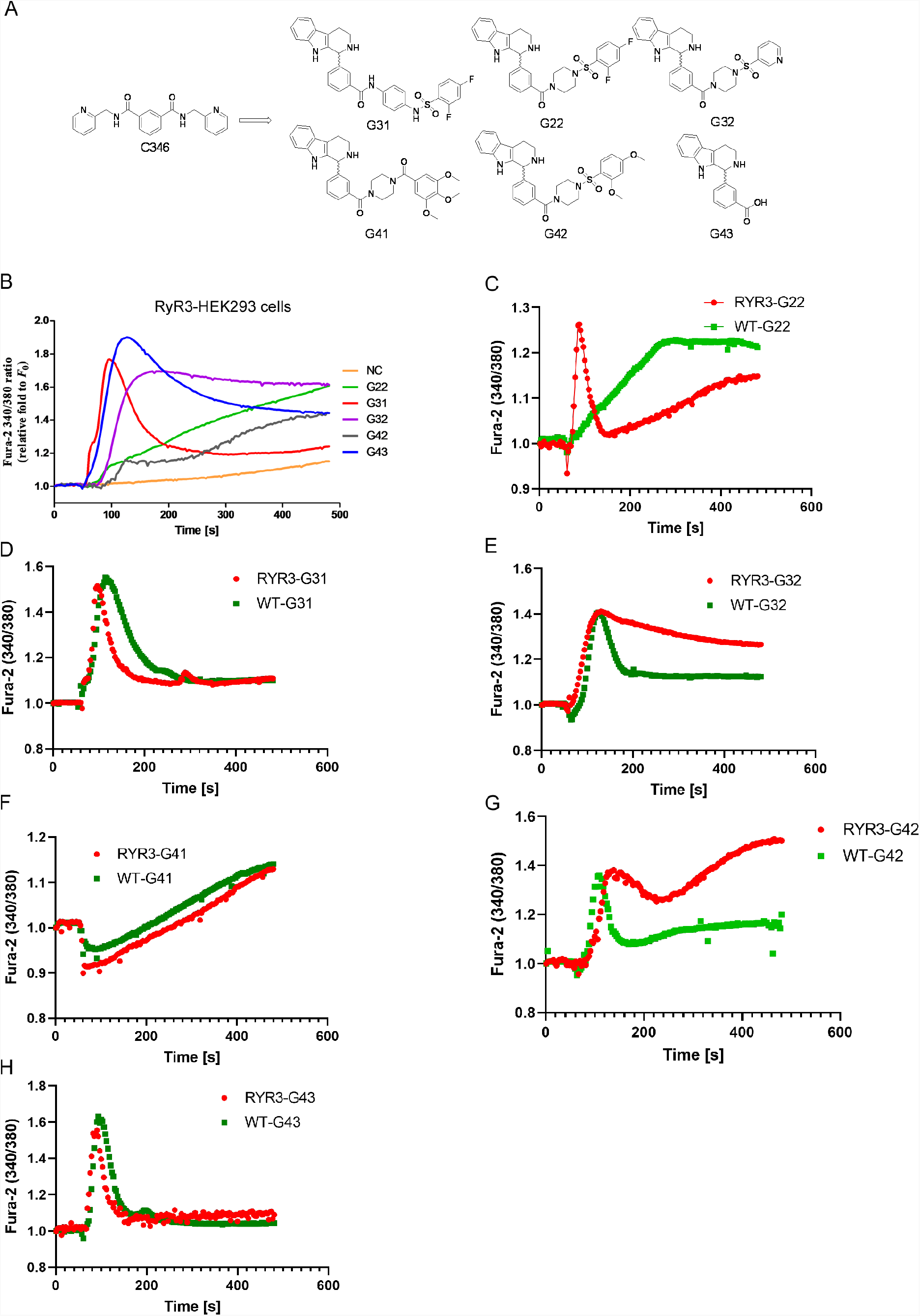
C346 analogs induce intracellular Ca^2+^ mobilization. (**A**) Structures of C346 and its analogs. (**B**) The ability of C346 analogs (50 μ M) to stimulate intracellular Ca^2+^ concentration in RYR3-HEK293 cells. (**C-H**) The ability of C346 analogs (20 μM), G22 (**C**), G31 (**D**), G32 (**E**), G41 (**F**), G42 (**G**), or G43 (**H**), to induce intracellular Ca^2+^ concentration in RYR3-HEK293 cells or control HEK293 cells.

We then set out to determine the source of the first Ca^2+^ peak triggered by G42. Interestingly, pretreatment of HEK293 cells with 2-APB, an inhibitor of IP3R and TRP channels (Ma et al., 2001; Prakriya and Lewis, 2001), significantly inhibited the first intracellular Ca^2+^ spike induced by G42 (**Fig. 4A**). In addition, we transfected HEK293T cells with GECO-TRPML1, a lysosomal-targeted Ca^2+^ sensor (Dong et al., 2010), and showed that G42 significantly induced lysosomal Ca^2+^ release (**Fig. 4B**), suggesting that G42 might target lysosomal Ca^2+^ channels. Since TRPMLs and TPCs are the two major lysosomal calcium channels, we first constructed the TRPML1^△NC^-expressing HEK293 cells, in which TRPML1 was expressed to the plasma membrane(Gerndt et al., 2020a). We showed that ML-SA1, a selective and potent agonist of TRPMLs (Shen et al., 2012), or G42 induced Ca^2+^ influx in TRPML1L15L/AA-L577L/AA-HEK293 cells (**Fig. 4C**). Likewise, we constructed TPC2L11A/L12A-expressing HEK293 cells, in which TPC2 was redirected to the plasma membrane (Gerndt et al., 2020b). G42 induced Ca^2+^ influx in TPC2L11A/L12A-expressing HEK293 cells (**Fig. 4D**). These results suggest that G42, in addition to its involvement in RYR-mediated intracellular Ca^2+^ mobilization (sustained or second Ca^2+^spike in **Fig. 3G**), also significantly induces lysosomal Ca^2+^ release (first Ca^2+^ spike in **Fig. 3G**).

**Figure 4.**
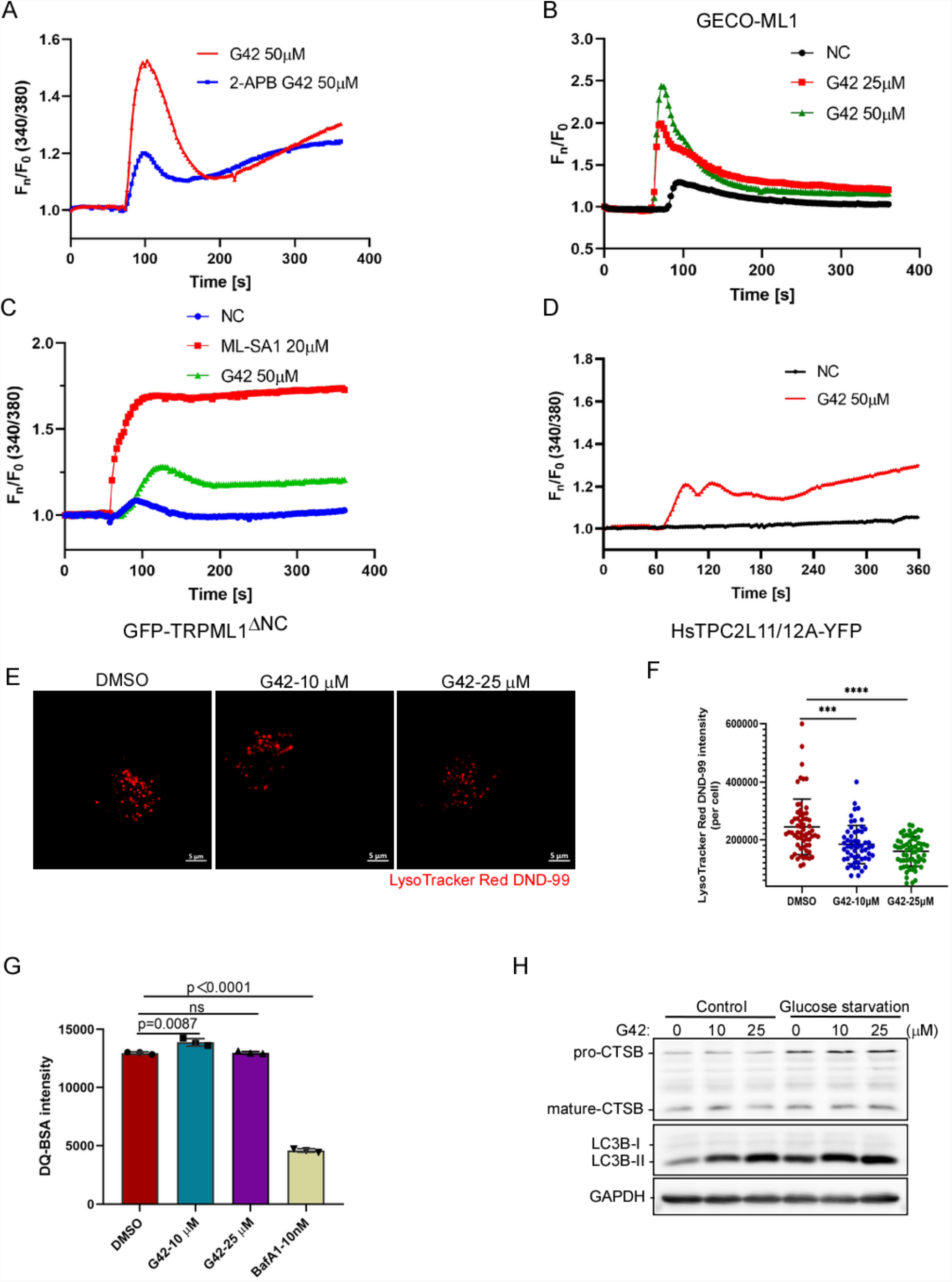
G42 significantly induces lysosomal Ca^2+^ release and alkalizes lysosomal pH. (**A**) HEK293 cells were pretreated with or without 2-APB (10 μM), the ability of G42 to trigger intracellular Ca^2+^ release was examined. (**B**) HEK293 cells were transfected with GECO-TRPML1; At 24 h post-transfection, cells were labeled with fura-2 and examined the ability of G42 to trigger lysosomal calcium release in GECO-ML1-overexpressing cells. **(C)** HEK293 cells were transfected with GFP-TRPML1^△NC^; At 24 h post-transfection, cells were labeled with fura-2 and examined the ability of G42 to trigger Ca^2+^ influx in GFP-TRPML1^△ NC^ positive cells. (**D**) G42 induced Ca^2+^ influx in TPC2L11A/L12A-expressing HEK293 cells. HEK293 cells were transfected with HsTPC2L11/12A-YFP; At 24 h post-transfection, cells were labeled with fura-2 and examined the ability of G42 to trigger Ca^2+^ influx in HsTPC2L11/12A-YFP-expressing cells. (**E, F**) G42 (10 or 25 μM) led to an increase of lysosomal pH in HeLa cells, manifested by the fluorescence intensity decreases of LysoTracker Red DND-99-labeled cells. (**G**) HeLa cells treated with or without the indicated compound were labeled with 10 μg/ml DQ-green BSA, followed by flow cytometry analysis. (**H**) HeLa cells were cultured in the presence or absence of glucose and were then treated with or without G42 at indicated concentration, followed by immunoblot analysis against CTSB, LC3II, and GAPDH.

### G42 inhibits autophagosome-lysosome fusion

Since lysosomal Ca^2+^ mobilization is coupled to lysosomal pH (Lu et al., 2013), we next assessed the effects of G42 on lysosomal pH. We treated the HeLa cells with or without G42, and then labeled them with LysoTracker Red DND-99. As expected, G42 did increase lysosomal pH in a concentration-dependent manner (**Figs 4E** and **4F**). Since alkalized lysosomes might compromise endolysosomal degradation, we tested the ability of G42 to inhibit endosomal trafficking of DQ-BSA. When DQ-BSA (a self-quenching dye) is delivered to the lysosome via endocytosis, its degradation relieves fluorescence-quenching and generates fluorescent cells (Marwaha and Sharma, 2017). To our surprise, G42 had little effect on the degradation of DQ-BSA, whereas Bafilomycin (Baf-A1), a vacuolar ATPase inhibitor, markedly inhibited its degradation (**Fig. 4G**). Likewise, G42 had little effect on the maturation of cathepsin B (CTSB) (**Fig. 4H**). These results suggest that G42-induced subtle alkalization of lysosomes does not change the lysosomal degradation function.

Since lysosomal Ca^2+^ mobilization is also involved in the fusion of autophagosomes and lysosomes (Medina, 2021), we examined the ability of G42 to modulate autophagy. We showed that G42 significantly increased the expression of LC3-II and p62 (**Fig. 5A**). The combination of G42 and Baf-A1, which is commonly used to block the autophagosome-lysosome fusion, did not further increase the Baf-A1-stimulated expression of LC3-II and p62, whereas G42 increased starvation-stimulated LC3-II and p62 expression (**Fig. 5B**). These data suggest that G42 is a late-stage autophagy inhibitor that blocks autophagosome-lysosome fusion, leading to the accumulation of LC3-II and p62. Also, in GFP-RFP-LC3-expressing HeLa cells, G42, like Baf-A1, significantly promoted the accumulation of yellow LC3 puncta (representing autophagosomes), whereas starvation stimulated both red-only LC3 puncta (representing autolysosomes) and yellow LC3 puncta (**Fig. 5C**). Moreover, in HeLa cells expressing RFP-LC3 and GFP-STX17, like Baf-A1, G42 markedly induced colocalization of LC3 red puncta with STX17 green puncta (**Fig. 5D**). In summary, these results indicate that G42 is a late-stage autophagy inhibitor.

**Figure 5.**
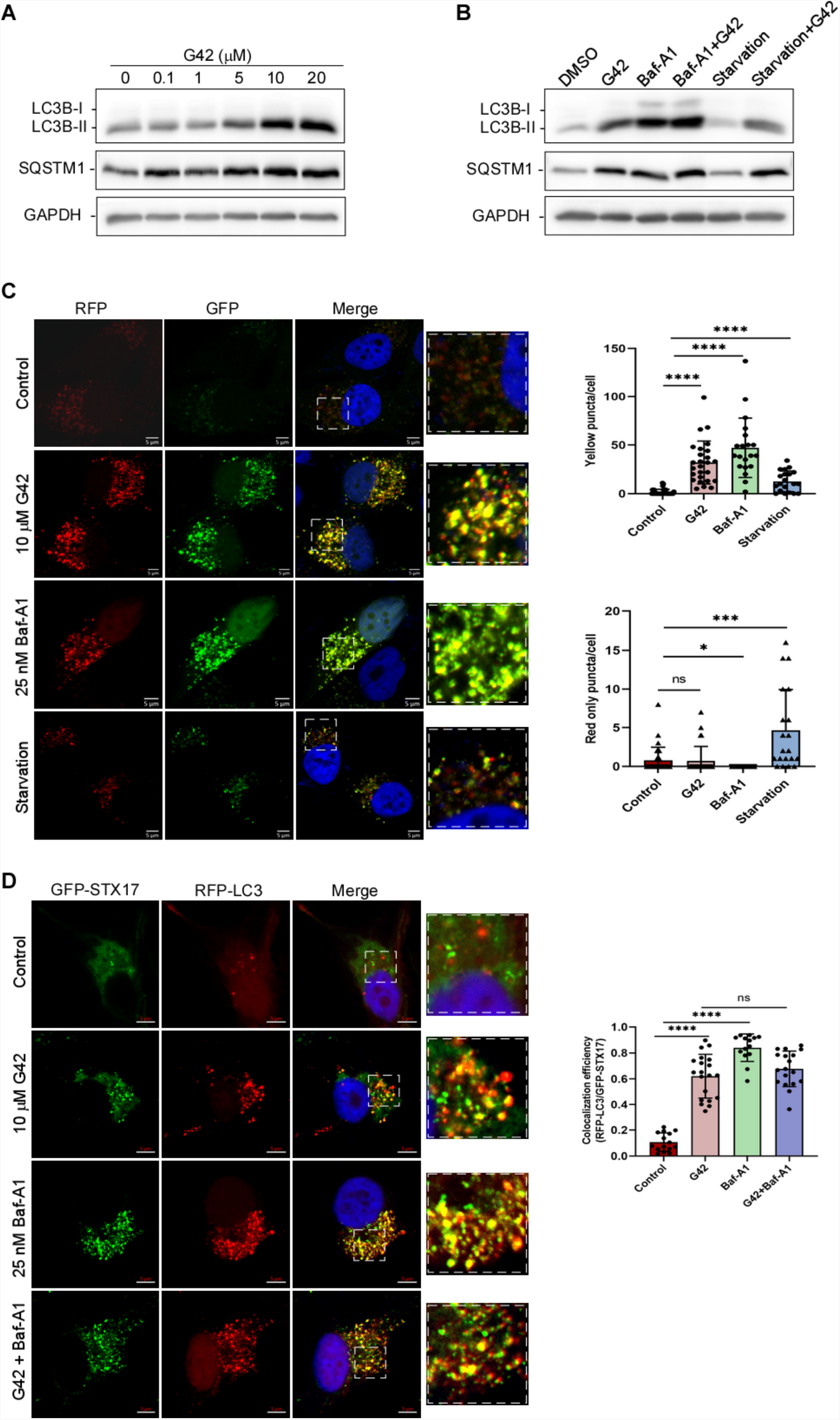
G42 is a late-stage inhibitor of autophagy. (**A**) G42 treatment of HeLa cells significantly increased the expression of LC3-II and p62. HeLa cells were treated with the indicated dose of G42 for 6 h; then cell lysates were collected and subjected to western blot analysis. (**B**) G42 is a late-stage autophagy inhibitor. HeLa cells were treated as indicated, followed by immunoblot analysis against CTSB, LC3II, and GAPDH. (**C**) G42 induced autophagosome accumulation. Under the treatment of G42, bafilomycinA1 (Baf-A1) or glucose starvation for 6 h, tf-LC3 stably expressing HeLa cells were captured by confocal imaging. (**D**) G42-stimulated LC3 red puncta colocalized with STX17 green puncta. HeLa cells that stably expressed RFP-LC3 and GFP-STX17 were treated with the indicated compound for 6 hours before being subjected to confocal imaging.

### G42 inhibits ZIKV and MHV infections of host cells

We previously showed that late-stage autophagy inhibitors, e.g., enanderinanin J, or saikosaponin D, potently inhibited Zika virus (ZIKV), Japanese encephalitis virus (JEV), or enterovirus A17 (EV-A17) infection of host cells (Huang et al., 2021b; Li et al., 2019). We, thus, examined whether G42 is a potential antiviral agent because of its autophagy inhibitory activity. We pretreated A549 cells with different concentrations of G42, followed by ZIKV infection. We showed that pretreatment of A549 cells with G42 significantly inhibited ZIKV-induced ZIKV envelope protein (ZIKV-E) (**Fig. 6A**). Likewise, pretreatment of A549 cells with G42 significantly suppressed ZIKV-induced double-stranded RNAs (dsRNAs), which are an intermediate in viral genome replication (**Figs 6B** and **6C**). The IC_50_ of G42 against ZIKE was around 1.33 μM (**Fig. 6C**). Pretreatment of A549 cells with G42 also significantly inhibited ZIKV-induced intracellular and extracellular virus titers (**Figs 6D** and **6E**). We last set out to determine whether G42 inhibits the entry, post-entry, or both of ZIKV infection. As shown in **Figure 6F**, A549 cells were treated with DMSO or G42 (25 μM) for the indicated time before or after ZIKV infection. Cells were incubated with 5 MOI of ZIKV for 1 h and then washed with PBS three times to remove the un-bound virion particles. Cells were incubated with a fresh medium that contained DMSO or G42 for an additional 17 h. We showed that pretreatment or posttreatment of G42 after viral entry all significantly inhibited ZIKV propagation, indicating that G42 affects different stages of the ZIKV life cycle. One possibility is that G42 might inhibit the early stage of ZIKV life cycle due to its alkalization effect on acidic compartments, e.g., late endosomes. Moreover, treatment of A549 cells after ZIKV infection with G42 inhibited ZIKV-induced intracellular and extracellular virus titers as potently as pretreatment of cells with G42 (**Figs 6F** and **6G**). These results suggest that G42 inhibits ZIKV infections after virus entry, likely by blocking autophagy.

**Figure 6.**
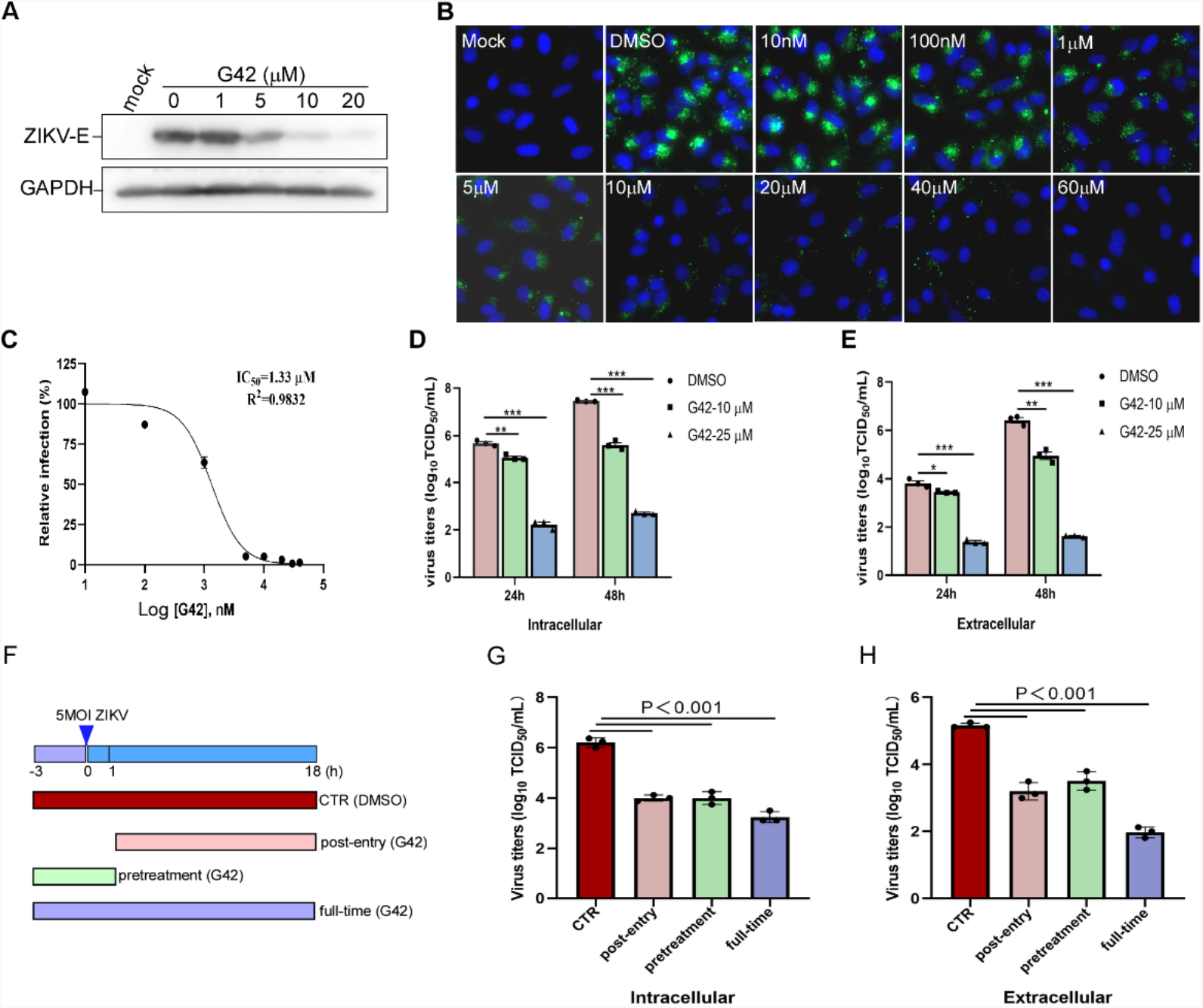
G42 effectively inhibits the infection of host cells by ZIKV. (**A**) G42 significantly inhibited the expression of ZIKV envelope protein (ZIKV-E) induced by ZIKV infection in A549 cells. A549 cells treated with the indicated concentration of G42 were infected with 0.5MOI of ZIKV; At 24 hours post-infection, cell lysates were collected, followed by immunoblot analysis against ZIKV envelope protein (ZIKV-E) and GAPDH. (**B**) G42 treatment of A549 cells significantly inhibited ZIKV infection. A549 cells were pretreated with the indicated concentration of G42 for 3 h, and then infected with 2 MOI of ZIKV. Cells were fixed and immunostained with dsRNA antibody. (**C**) Dose-response relationship between G42 and ZIKV infectivity. A549 cells were pretreated with the indicated concentration of G42 for 3 h, and then infected with 0.5 MOI of ZIKV; Quantitative PCR (qPCR) was performed to analyze the level of the intracellular viral genome. (**D, E**) G42 pretreatment of A549 cells significantly inhibited the production of intracellular (**D**) and extracellular (**E**) viral particles after ZIKV infection. (**F-H**) A549 cells were treated with or without G42 for the indicated time course before or after ZIKV infection (**F**). Intracellular (**G**) and extracellular (**H**) viral particles after ZIKV infection were then determined.

Recently, chloroquine, a late-stage autophagy inhibitor similar to G42, exhibited anti-SARS-CoV-2 activity (Wang et al., 2020). We, therefore, assessed whether G42 inhibits murine hepatitis virus (MHV) (Bosch et al., 2004), a β-coronavirus, infection of its host cells. We showed that treatment of 17-Cl1 cells with G42 inhibited the MHV-induced NSP9 expression dose-dependently (**Figs 7A**). Likewise, G42 efficiently inhibited the expression of NSP9 induced by different MOI MHV (**Fig. 7B**). Moreover, G42 significantly inhibited MHV-induced dsRNA levels in 17-Cl1 cells with IC_50_ around 1.4 μM (**Fig. 7C**). These results suggest that G42 is an anti-β-coronavirus agent.

**Figure 7.**
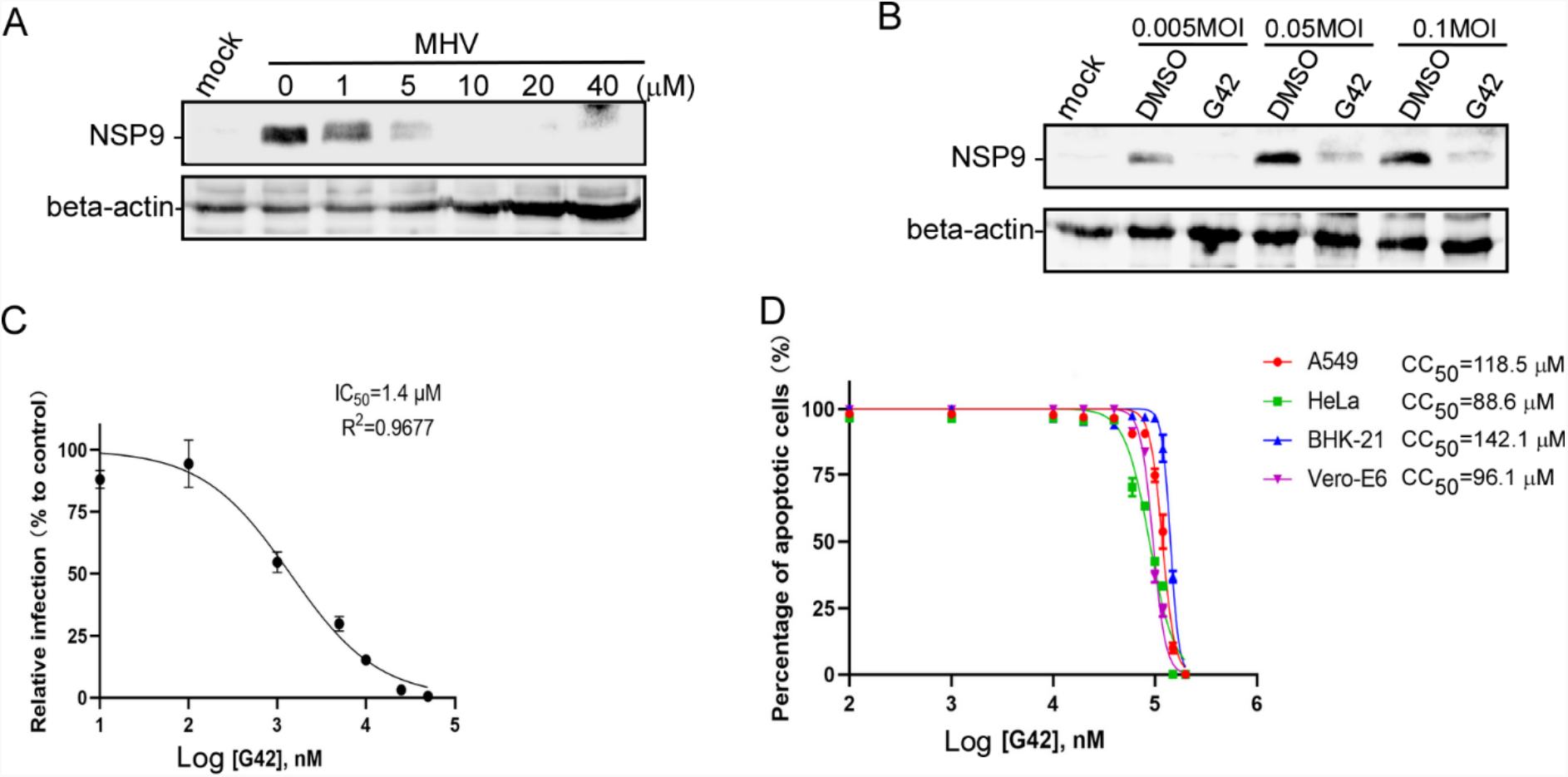
G42 effectively inhibits the infection of host cells by MHV and exhibits low cytotoxicity. **(A)** Western-blot analysis of G42 treated 17-Cl1 cells at 18 hours after MHV infection. 17-Cl1 cells pretreated with indicated dose of G42 for 3 hours were infected with 0.05 MOI of MHV; At 1 hour post-infection, the medium containing viruses was removed, cells were washed with PBS and incubated with a fresh medium containing an indicated dose of G42. (**B**) G42 treatment of 17-Cl1 cells significantly inhibited the expression of NSP9 induced by different MOI MHV infections. (**C**) G42 treatment of 17-Cl1 cells significantly inhibited the dsRNA-positive structure induced by MHV infection. (**D**) The cytotoxicity of G42 in different cell lines.

Manipulating the intracellular Ca^2+^ levels or Ca^2+^ signals has been shown to inhibit the virus infection (Alam et al., 2016; Fujioka et al., 2018; Sakurai et al., 2015; Scherbik and Brinton, 2010; Wang et al., 2017). The cytosolic Ca^2+^ levels or related signaling are essential to cell viability; blockade of extracellular Ca^2+^ influx or release of Ca^2+^ from the ER or mitochondria is detrimental to cells. Therefore, using general Ca^2+^ channels or signaling inhibitors as antiviral agents is impractical. Here we showed that G42 triggered Ca^2+^ release from lysosomes and exhibited potent anti-ZIKV and anti-MHV activity, with IC_50_ around 1 μM, whereas its cytotoxicity was low (CC_50_ around 88-142 μM in different cell lines) (**Fig. 7D**). It is of interest to further determine the effects of G42 against other positive-sense single-stranded RNA viruses ((+)ss RNA viruses), e.g., SARS-CoV, MERS-CoV, JEV, and Dengue virus (DENV). In addition, modifying G42 structure might further improve its antiviral activity while decreasing its cytotoxicity. It also remains to determine the lysosomal Ca^2+^ channels targeted by G42, and the potential targets include TRPMLs (Venkatachalam et al., 2015; Wang et al., 2014), TPCs (Grimm et al., 2017; Patel, 2015), or both.

We previously showed that knockdown of ATG5 or ATG7, the genes essential for autophagy, inhibited ZIKV, JEV, or EV-A71 infection of cells (Huang et al., 2021b; Li et al., 2019). Also, treatment of cells with autophagy inhibitors, e.g., enanderinanin J, saikosaponin D, chloroquine, or G42 inhibited various (+)ss RNA viruses, e.g., ZIKV, JEV, EV-A71, and MEV (Huang et al., 2021b; Li et al., 2019; Wang et al., 2020). Since these inhibitors inhibited the autophagosome-lysosome fusion, these RNA viruses might hijack autolysosomes as replication sites. Alternatively, it is possible that the autolysosomes might be the site for the degradation of the synthesized negative-strand viral RNA, and in this way, it facilitates the delivery of the replicated positive-strand viral RNA to ER for viral protein synthesis. Yet, the role of autophagy in viral infection is complicated, depending upon the viral types, host cells, infection duration, and stages of viral cycles. Thus, more efforts are needed to systematically dissect the role of autophagy in viral infections.

## MATERIALS AND METHODS

### Chemical Synthesis of C346 analogs

The synthesis of the analogs of C346 was illustrated in Schemes 1 – 4 as follows.

#### 2,4-difluoro-N-(4-nitrophenyl)benzenesulfonamide (2)

4-Nitrobenzenamine (**1**) and pyridine (1.2 eq) were dissolved in DCM, to which 2,4-difluorobenzenesulfonyl chloride (1.05 eq) was added in portion. The mixture was stirred at room temperature for 16 hours before quenched with water. The organic layer was separated, washed with brine, dried over anhydrous Na_2_SO_4,_ and concentrated under a vacuum. The residue was used directly in the next step without further purification.

#### N-(4-aminophenyl)-2,4-difluorobenzenesulfonamide (3)

Compound **2** was dissolved in THF/EtOH/H_2_O, to which Fe powder (5.0 eq) and NH_4_Cl (10 eq) were added. The resulting mixture was refluxed for 2 hours and filtered with celite to remove the insoluble solid. The volatiles were removed under reduced pressure. The residue was then resolved in EA, washed with water and brine, dried over anhydrous Na_2_SO_4_, and concentrated under a vacuum. The obtained residue was used directly in the next step without further purification.

#### N-(4-((2,4-difluorophenyl)sulfonamido)phenyl)-3-formylbenzamide (4)

Compound **3** was dissolved in DCM, to which 3-carboxybenzaldehyde (1.05 eq) and Et_3_N (3.0 eq) were added. Propylphosphonic anhydride (T3P, 1.1 eq) was then added dropwise into the mixture, which was stirred at room temperature for 16 hours. The mixture was quenched with water before the organic layer was separated. The organic layer was washed with water and brine, dried over anhydrous Na_2_SO_4,_ and concentrated under a vacuum. The residue was used directly in the next step without further purification.

#### N-(4-((2,4-difluorophenyl)sulfonamido)phenyl)-3-(2,3,4,9-tetrahydro-1H-pyrido[3,4-b]indol-1-yl)benzamide (**G31**)

Compound **4** was dissolved in EtOH with AcOH (catalytic amount). Tryptamine (1.0 eq) was then added to the mixture, which was refluxed for 16 hours. The volatiles were removed under a vacuum. The residue was purified with column chromatography (silica gel, DCM/MeOH) to afford compound **G31**.

*Compounds* ***6a*-6c** were prepared from 1-boc-piperazine (**5**) and ArSO_2_Cl with the same synthetic method of compound **2**.

#### Compounds 7a –7c

The corresponding compound **6** was dissolved in DCM, to which trifluoroacetic acid (10 eq) was added. The mixture was stirred at room temperature for 1 hour and concentrated under a vacuum. The residue was used for the next step without further purification.

Compounds **8a** – **8c** were prepared from the corresponding **7a** – **7c** with the same synthetic method of compound **4**.

Compounds **G22, G32**, and **G42** were prepared from corresponding **8a** – **8c** with the same synthetic method of compound **G31**.

#### tert-butyl 4-(3,4,5-trimethoxybenzoyl)piperazine-1-carboxylate (9)

1-boc-piperazine (**5**) was dissolved in DCM, to which pyridine (2.0 eq) and 3,4,5-trimethoxybenzoylchloride (1.0 eq) were added dropwise. The mixture was stirred at room temperature for 16 hours and washed with water and brine. The organic layer was then dried over anhydrous Na_2_SO_4_ and concentrated under a vacuum. The residue was used directly in the next step without further purification.

Compounds **10, 11**, and **G41** were prepared with the same synthetic methods as compounds **7, 4**, and **G31**, respectively.

**Scheme 1.**
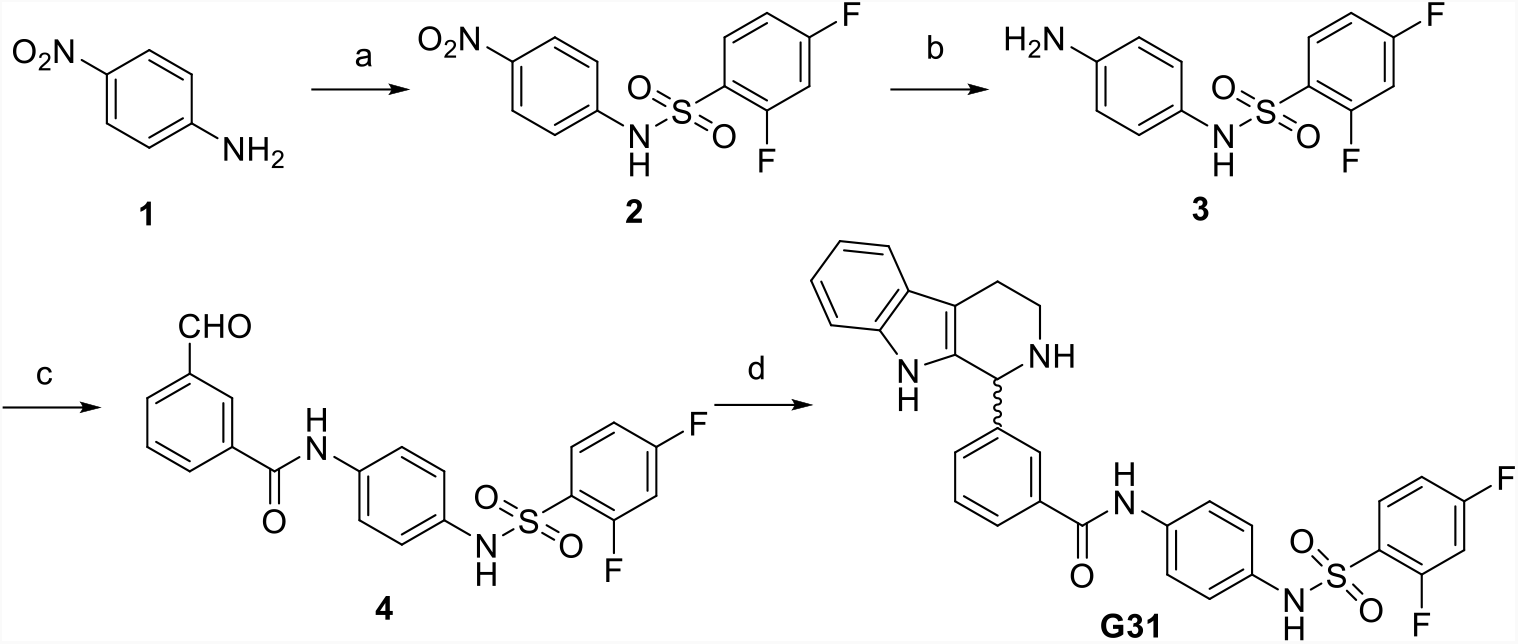
Synthesis of compound **G31**. Reagents and conditions: (a) 2,4-difluorobenzenesulfonyl chloride, pyridine, DCM, rt; (b) Fe, NH_4_Cl, THF/EtOH/H_2_O, reflux; (c) 3-carboxybenzaldehyde, T3P, Et_3_N, DCM, rt; (d) tryptamine, AcOH, EtOH, reflux.

**Scheme 2.**
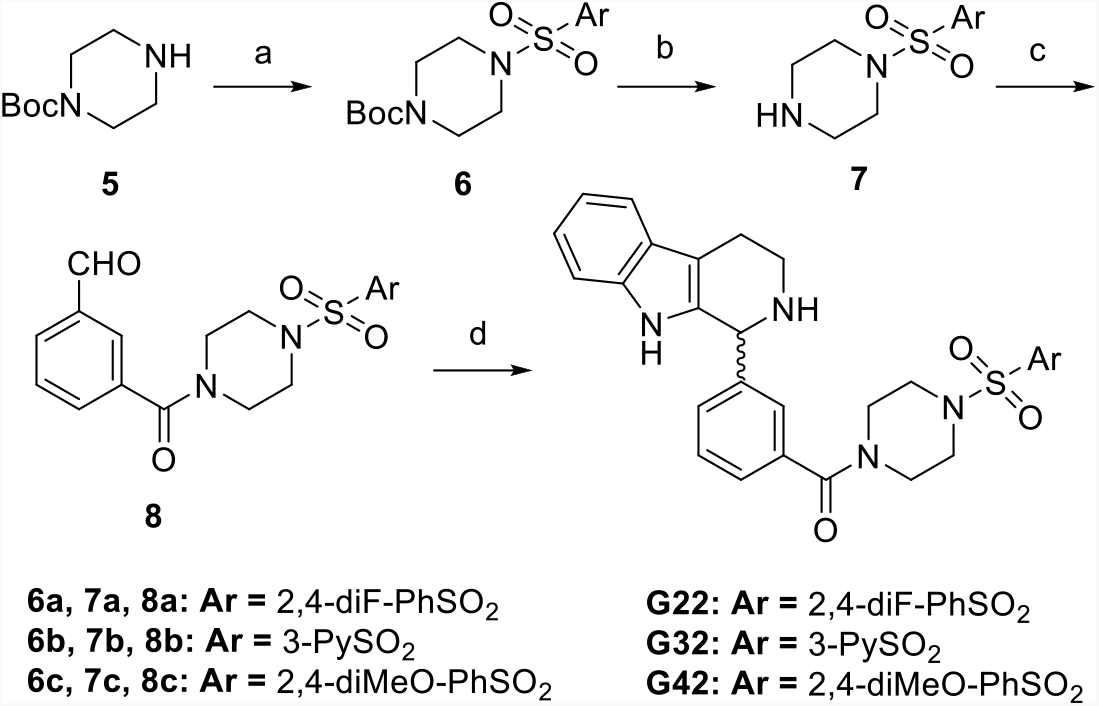
Synthesis of compounds **G22, G32**, and **G42**. Reagents and conditions: (a) ArSO_2_Cl, pyridine, DCM, rt; (b) TFA, DCM, rt; (c) 3-carboxybenzaldehyde, T3P, Et_3_N, DCM, rt; (d) tryptamine, AcOH, EtOH, reflux.

**Scheme 3.**
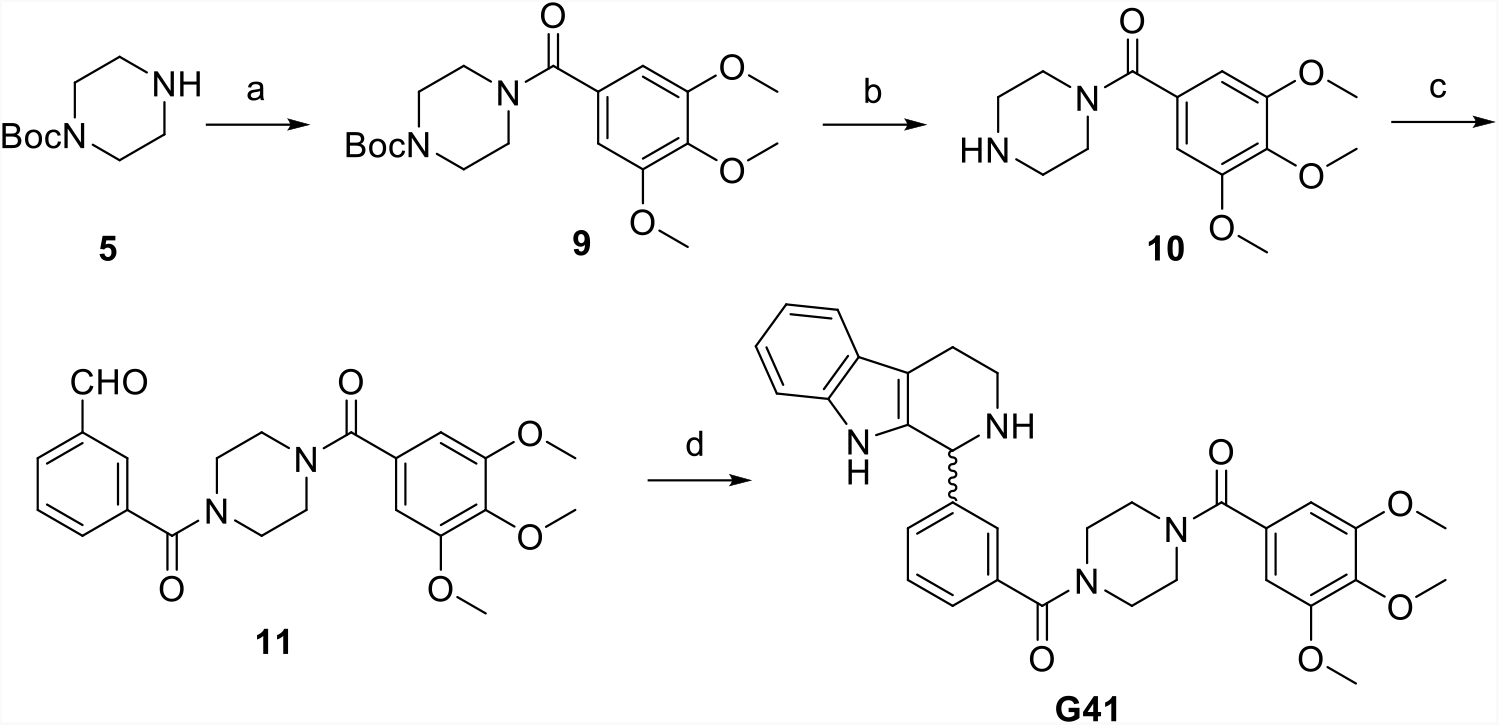
Synthesis of compound **G41**. Reagents and conditions: (a) 3,4,5-trimethoxybenzoylchloride, pyridine, DCM, rt; (b) TFA, DCM, rt; (c) 3-carboxybenzaldehyde, T3P, Et_3_N, DCM, rt; (d) tryptamine, AcOH, EtOH, reflux.

**Scheme 4.**
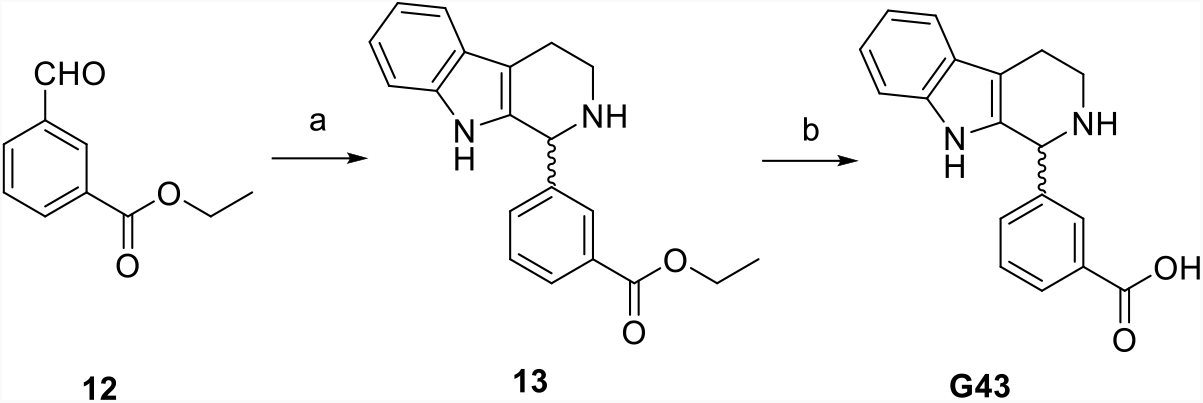
Synthesis of compound **G43**. Reagents and conditions: (a) tryptamine, AcOH, EtOH, reflux; (b) LiOH, THF/EtOH/H_2_O, rt. Compound **13** was prepared with the same method as compound **G31** but without further purification.

#### 3-(2,3,4,9-tetrahydro-1H-pyrido[3,4-b]indol-1-yl)benzoic acid (G43)

Compound **13** was dissolved in THF/EtOH/H_2_O, to which LiOH (10 eq) was added. The mixture was stirred at room temperature for 16 hours and concentrated under a vacuum. The residue was purified with column chromatography (silica gel, DCM/MeOH) to afford compound **G43**.

### Virtual screening

The virtual screening was performed as described previously ^6^.

### Materials for chemical synthesis

All starting materials, reagents, and solvents were purchased from commercial suppliers and directly used in the synthesis without further purification.

### Cell culture

Human cervical adenocarcinoma cells (HeLa), human lung carcinoma cells (A549), African green monkey kidney cells (Vero-E6), Baby hamster kidney cells (BHK-21), and murine 17Cl-1 cells were cultured in Dulbecco’s Modified Eagle’s Medium (DMEM) supplemented with 10% FBS (Gibco, 10500064) and 100 U/mL of penicillin-streptomycin (Gibco, 15140122).

### Viruses

The ZIKV PRVABC59 stain (ATCC, VR-1843) was propagated in A549 cells. The MHV A59 stain (ATCC, VR-764) was cultured in 17Cl-1 cells for amplification.

### Plasmids

The pLenti-RFP-tagged, or mRFP-GFP tandem tagged MAP1LC3 (tf-LC3) construct was constructed by our lab as previously described (Wang et al., 2016). TPML1 (ML1)-GECO was a gift from Dong Xianping. HsTPC2L11/12A-YFP and TRPML1ΔNC-GFP were generously provided by Sandip Patel.

### Intracellular Ca^2+^ measurements

Control or Ryanodine receptor 3 (RYR3)-overexpressing HEK293 cells were plated in 24-well plates. The next day, the cells were incubated with 4 μM Fura-2 AM (Invitrogen, F1221) and 0.4% Pluronic™ F-127 (Invitrogen, P3000MP) in HBSS (Gibco, 14025092) at room temperature (RT) for 30 min. After being washed with Ca^2+^-free HBSS containing 2 mM EGTA, the cells were incubated in Ca^2+^-free HBSS with or without indicated compounds at RT for 30 min. Fluorescence images were acquired at 3 s intervals by alternate excitation at 340 nm and 380 nm with emission at 510 nm in a Nikon Eclipse Ti-S Calcium imaging system.

To measure the lysosomal Ca^2+^ release, ML1-GECO was transfected to HEK293T cells. At 24 h post-infection, the fluorescence intensity at 470 nm was monitored using the Nikon Eclipse Ti-S Calcium imaging system. Lysosomal Ca^2+^ release was measured under Ca^2+^-free HBSS containing 2 mM EGTA.

### Western blot analysis

Cells were treated indicated dosage of G42, and total protein samples were then harvested by using cell lysis buffer (Beyotime, P0013J). The protein concentration of cell lysates was determined by the Bradford assay (Bio-Rad Laboratories). An equal amount of protein samples was separated using SDS-PAGE gel electrophoresis. The proteins were then transferred to a PVDF membrane (Millipore), blocked with 5% non-fat milk, and blotted sequentially with primary and secondary antibodies.

### Lentivirus production and stable cell line generation

T1×10^6^ HEK293T cells were plated in gelatin-coated 6-well plates. Next day, transfections were performed using the the pLenti-CMV-DEST vectors together with the lentivirus envelope and package plasmids pMD2.G (Addgene, #12259) and psPAX2 (Addgene, #12260). The transfection mixture containing three plasmids was incubated at RT for 5 min, and then PEI was added at a 1:3 ratio. After incubation at RT for 10 min, the mixture was added to the cells. Four hours post-transfection, the transfection medium was removed and changed with a fresh complete medium. The viruses were collected every 24 hours and stocked at -80°C.

### Viral titer measurement

The virus titer of viral samples was determined by TCID_50_ assay as previously described(Huang et al., 2021a).

### DQ-Green BSA trafficking assay

HeLa cells were plated on the 6-well plates. The next day, cells were treated with or without bafilomycin A1 (10 nM) or G42 (10 μM or 25 μM) containing 10 μg/mL DQ-Green BSA (Invitrogen, D12050) at 37°C for 6 h followed by FACS analysis.

### Lysotracker pH measurement

HeLa cells were treated with indicated dose of G42 (DMSO as control) for 5 h, and then changed medium containing 100 nM Lysotracker Red DNS-99 and indicated dose of G42 at 37°C for 1 h. Cells were washed twice with PBS and changed with fresh DMEM. Images were captured by the LSM780 confocal microscope (Carl Zeiss USA). The fluorescence intensity of each cell was quantified on Zeiss Zen 2 Offline Station.

### Immunofluorescence staining

Cells were fixed with 4% paraformaldehyde (PFA) at room temperature for 15 min. After being rinsed with PBS, cells were incubated with PBS containing 5% normal donkey serum (Chemicon, S30-100ML) and 0.2% Triton X-100 at RT for 1 h, and then blocked with an anti-dsRNA antibody (Scicons, 10010200) and a fluorescent secondary antibody (Invitrogen). Images were captured with the CellInsight CX7 High-Content Screening (HCS) Platform (Thermo Scientific).

### Cytotoxicity analysis

.BHK-21, A549, HeLa, or Vero-E6 cells were seeded in 96-well plates (Corning, 3603) and incubated with or without G42 at the indicated concentration. Live cells were then stained with propidium iodide (Invitrogen, P3566) and Hoechst 33258 (Invitrogen, H3570). Images were finally captured with the CellInsight CX7 High-Content Screening platform with a 10× objective lens. The dead cells in each group were quantified with an HCS Studio ™ 3.0 (Thermo Fisher). The half-maximal cytotoxic concentrations (CC50) were analyzed with Graphpad Prism 9.

### Statistical analysis

Data were presented as mean ± SD. Statistical analysis was calculated by GraphPad Prism 9 software. An unpaired two-tailed student’s t-test was used to analyze the statistical significance of two groups. p < 0.05 was considered statistically significant. P > 0.05 ; *, P ⩽ 0.05 ; **, P ⩽ 0.01 ; ***, P ⩽ 0.001; ****, P ⩽ 0.0001.

## ACKNOWLEDGMENTS

We thank members of Yue laboratory for advice on the manuscript. This work was supported by Hong Kong Research Grant Council grants (11101717 and 11103620), NSFC (21778045, 32070702, and 82161128014), ITF (MRP/064/21, GHP/097/20GD, MHP/072/21), Shenzhen Science and Technology Innovation Committee (SGDX20201103093201010, JSGG20200225150702770, and JCYJ20210324134007020) and the Non-profit Central Research Institute Fund of Chinese Academy of Medical Sciences (2021-JKCS-013).

